# Diffusion-ACP39: A Decoder-Adaptive Latent Diffusion Framework for Generative Anticancer Peptide Discovery

**DOI:** 10.64898/2026.03.04.709539

**Authors:** Jielu Yan, Qichun Wu, Yifan Li, Jianxiu Cai, Mingliang Zhou, François-Xavier CACPbell-Valois, Shirley W. I. Siu

**Author notes:** These authors contributed equally to this work (Jielu Yan, Qichun Wu).

## Abstract

Cancer remains a major global health threat, with its incidence and mortality rates consistently rising in recent years. Anticancer peptides (ACPs) are short amino acid chains that can inhibit the growth or spread of cancer cells. Compared to traditional treatments, ACPs are a promising class of potential cancer therapies due to their multiple mechanisms, potential for combination cancer therapy, enhanced immune function, lower toxicity to normal tissues, fewer side effects, and less drug resistance. Although it is necessary to explore novel ACPs, traditional wet-lab methods for selecting them are labor-intensive, time-consuming, and expensive. To accelerate the discovery of novel ACPs, we proposed Diffusion-ACP39, a latent diffusion-based generative model with synchronized seed autoencoder for anticancer peptide design, capable of generating novel peptides with lengths ranging from 5 to 39 amino acids. Furthermore, we developed RF-ACP39, a random forest classifier model to assess the generative power of Diffusion-ACP39. Finally, Diffusion-ACP39 achieved an accuracy of 94.5% when generating 10,000 peptides with RF-ACP39. We also qualitatively analyzed the differences among true ACPs, random sequences, random peptides, and generated ACPs, demonstrating that the generated ACPs are most similar to true ACPs.

## 1 Introduction

Cancer remains one of the leading causes of global mortality. According to recent investigations, approximately 20 million new cancer cases and nearly 9.7 million cancer-related deaths occurred worldwide in 2022 [1]. The most common malignancies, ranked by projected incidence for 2025 in descending order, include breast cancer, prostate cancer, and lung and bronchus cancer. In the United States, the annual age-adjusted cancer incidence rate was 455.6 per 100,000 individuals during 2017-2021, with a mortality rate of 146 per 100,000 from 2018 to 2022 [2]. Epidemiologically, approximately 38.9% of individuals will receive a cancer diagnosis during their lifetime [1].

Conventional cancer interventions encompass surgery, chemotherapy, radiation therapy, targeted therapy, and immunotherapy. However, these approaches exhibit significant limitations. Surgery is inherently invasive and effective only for localized tumors, cannot eliminate micrometastases, and frequently necessitates adjuvant therapies. Chemotherapy suffers from high systemic toxicity, narrow therapeutic indices, and frequent development of drug resistance. Radiation therapy often causes collateral damage to adjacent healthy tissues, demonstrates limited efficacy against metastatic disease, and may induce tumor recurrence through selection of radiation-resistant clones. Targeted therapies are constrained by their dependency on specific molecular alterations, rapid emergence of resistance mechanisms, and high costs. Immunotherapies carry risks of severe immune-related adverse events, exhibit low response rates in immunologically “cold” tumors such as pancreatic carcinoma, and may paradoxically accelerate tumor progression in some cases.

Anticancer peptides (ACPs) represent a promising class of therapeutic agents characterized by short amino acid sequences that inhibit neoplastic proliferation and metastasis. These peptides employ multimodal mechanisms of action—including disruption of cancer cell membranes, angiogenesis inhibition, apoptosis induction, and immune response modulation—which collectively reduce the likelihood of resistance development compared to single-target agents. ACPs offer distinct advantages such as high tumor selectivity, low systemic toxicity, synergistic potential with existing therapies, ability to overcome conventional drug resistance pathways, and capacity to activate antitumor immunity. Furthermore, their modular structure permits rational design and optimization through in silico approaches, enabling enhancement of proteolytic stability, tissue penetration efficiency, and tumor-targeting specificity for novel therapeutic development.

To the best of our knowledge, only a few generation methods have been developed for novel ACPs. So far, Recurrent Neural Networks (RNNs) and a specific variant of Long Short-Term Memory (LSTM) networks, have been employed for generating novel ACP sequences. Lu *et al.* proposed a generative LSTM model trained on 584 ACP peptides, generating 40 novel ACP sequences, 75% of which achieved an ACP score higher than 90% when evaluated by an SVM classifier [3]. Zakharova *et al.* proposed a generative transfer RNN model that was fine-tuned on only 53 ACPs targeting HeLa cells, after being pre-trained on 4774 ACPs [4]. The fine-tuned model generated 50,000 novel ACP sequences, with approximately 20% predicted as potentially active by an RNN classifier. Ibrahim *et al.* proposed a generative RNN model trained on the acp740 dataset, followed by a three-tier filtration system to select high-potential and exclude less relevant ACP sequences [5]. The first tier consists of three classifiers that simultaneously identify high-potential ACPs. The second and third tiers aim to screen out less relevant sequences and perform a final filtration using a nearest centroid classifier and an unsupervised nearest neighbor learning approach, respectively. The generative RNN model achieved a combined hit-rate of 36.25% when generating 5000 novel ACP sequences as evaluated by the three classifiers in the first tier.

However, all existing generative methods for ACPs are based primarily on RNNs and their variant architectures, such as LSTMs [6, 7]. Compared to modern diffusion-based generative networks, these recurrent models exhibit several limitations. They often lack output diversity, tend to suffer from mode collapse, and are relatively time-consuming when generating novel sequences. Furthermore, RNNs and LSTMs struggle with capturing long-range dependencies and global context efficiently, which can restrict their ability to produce highly varied and structurally complex peptides. In contrast, diffusion models offer better parallelism, higher sACPle diversity, and more stable training dynamics [8].

Therefore, we propose Diffusion-ACP39, a latent diffusion-based generative model with synchronized seed autoencode, trained on an ACP dataset comprising 3490 sACPles with lengths ranging from 5 to 39, to generate novel ACPs within the same length range. Furthermore, a non-ACP dataset of equal size and similar length distribution was constructed as a negative set to train the classifier models. Diffusion-ACP39 is predicated on a novel “Synchronized Seed Autoencoding” mechanism. Deviating from conventional paradigms that rely on pre-trained, static latent spaces, we introduce a “Generation-First, Decoder-Adaptive” training strategy. Rather than performing generation within a fixed manifold, we initially train the diffusion model utilizing random projections anchored by fixed seeds, thereby empowering the U-Net to autonomously learn and define the data distribution within the latent space. Subsequently, the decoder is trained using identical seeds, enabling it to specifically adapt to and precisely interpret the unique latent feature representations yielded by the U-Net. This paradigm delegates the construction of latent semantics to the U-Net, while the decoder is tasked with materializing this space into discrete sequences, culminating in a logically self-consistent and highly efficient generative system. Finally, multiple classifier models were trained on the ACP and non-ACP datasets using the Random Forest (RF) algorithm. The model demonstrating the superior classification performance was selected to assess the quality and effectiveness of the generated sequences.

## 2 Material and Methods

### 2.1 Data collection

We collected an ACP dataset to develop our generative model, aiming to provide high-quality biological sequence features. All data underwent screening and preprocessing to ensure uniqueness and validity, resulting in a unified and core ACP experimental dataset. This initial dataset comprised 3,671 unique anticancer peptide sequences. Statistical analysis of sequence lengths revealed a broad range from 5 to 321 amino acid residues, fully reflecting the structural diversity of ACPs. The dataset was de-duplicated, with each sample containing a unique identifier, amino acid sequence, and corresponding length information, serving as the foundational data source for model training and analysis in this study.

#### 2.1.1 ACP dataset

After screening and preprocessing the collected ACP dataset, we formed a new positive sample dataset, ultimately retaining 3,489 high-quality sequences. To balance model training efficiency with the typical characteristics of bioactive peptides, sequence lengths were strictly limited to a range of 5 to 39 amino acid residues. We selected this length range because sequences between 5 and 39 residues account for nearly 95% of the baseline dataset, which is sufficient to encompass the majority of sequence characteristics.Outliers exceeding this length were excluded to ensure the consistency and stability of the model’s input distribution.

#### 2.1.2 Negative dataset

To ensure a balanced training dataset and mitigate potential bias arising from sequence length, we employed a random generation strategy based on length distribution matching to construct the non-ACP dataset. We strictly adhered to a length distribution identical to that of the positive ACP samples: for every sequence in the positive set, a corresponding random peptide of the same length was generated. During generation, each residue position was sampled uniformly from the 20 standard amino acids. Furthermore, to guarantee data validity and label purity, we implemented a rigorous exclusion mechanism: any generated sequence found in the positive ACP set or duplicated within the negative set was discarded and regenerated. This process yielded a final non-ACP dataset of 3,489 sequences, which preserves the exact length distribution of the positive set while exhibiting random and uniform amino acid composition.

Furthermore, we conducted a comparative analysis of the ACP and non-ACP datasets, covering sequence length distribution and amino acid composition characteristics. As shown in Figure 1, in terms of length distribution, the ACP dataset exhibited a high degree of consistency with the non-ACP dataset, with both having an average length of approximately 17.6, and the distribution curves almost completely overlapping. This indicates that the negative samples were strictly matched in length. However, significant differences were observed in amino acid composition. Compared to the non-ACP dataset, the ACP dataset was notably enriched in “K”, “L”, and “R”, while the contents of “M”, “E”, and “D” were relatively lower. This enrichment of positively charged amino acids aligns with the biophysical mechanism by which antimicrobial peptides typically adsorb onto bacterial cell membranes through electrostatic interactions.

**Figure 1:**
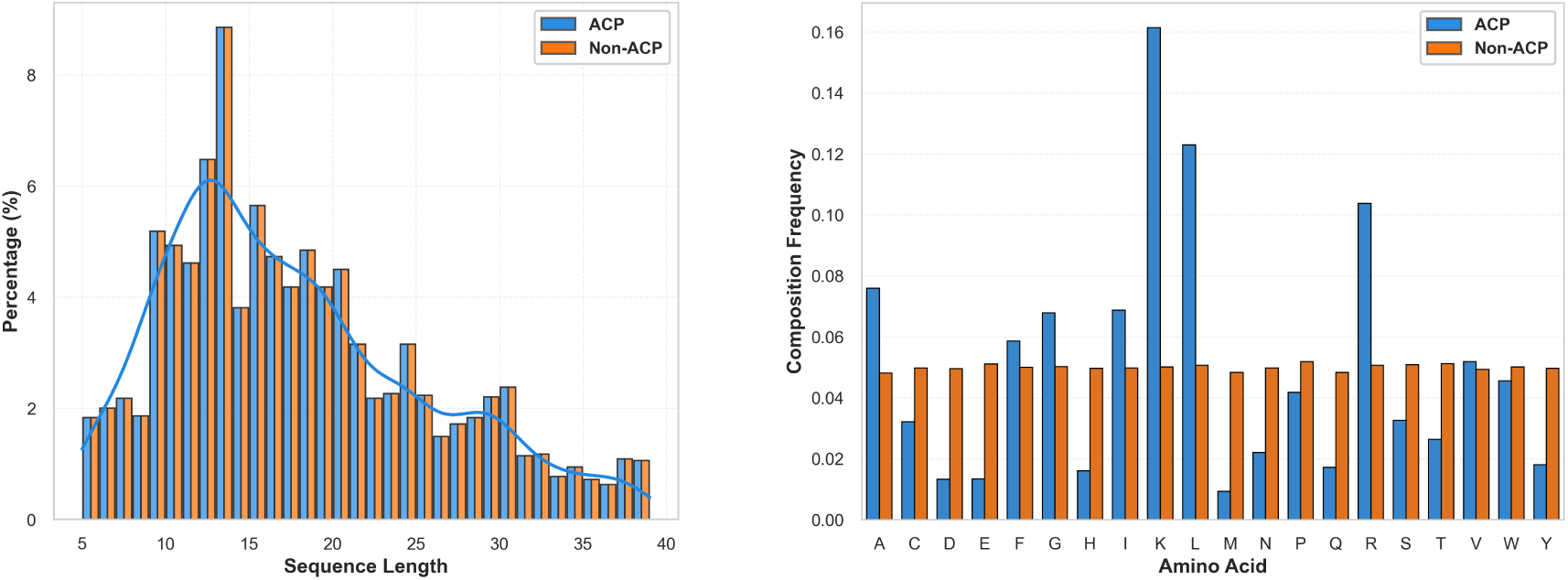
Comparison of length distribution (left) and amino acid composition (right) between ACP and non-ACP datasets.

### 2.2 Methods

#### 2.2.1 Performance metrics

We employed a comprehensive set of multi-dimensional performance metrics to evaluate the classifier models. Specifically, we reported test accuracy (Acc) as the baseline evaluation criterion. Considering the potential limitations of relying solely on accuracy for biological sequence data, we further incorporated the F1-Score and Matthews Correlation Coefficient (MCC). Notably, MCC is regarded as the most balanced metric for binary classification problems, as it accurately reflects the model’s predictive capability even when sample sizes are imbalanced. Additionally, the Area Under the Receiver Operating Characteristic Curve (ROC-AUC) was used to assess the model’s generalization ability across different thresholds. Finally, all experimental results were derived using 5-fold cross-validation to ensure statistical reliability.

#### 2.2.2 Feature Encoding methods

To comprehensively characterize protein sequences, we converted raw sequences into numerical feature vectors. First, we employed Amino Acid Composition (AAC) as the baseline feature. AAC captures global compositional information by calculating the frequencies of the 20 amino acids. To compensate for the lack of sequence order information in AAC, we introduced DDE and CKSAAP. DDE extracts local adjacency associations by calculating the deviation of dipeptides from their expected values, while CKSAAP focuses on residue pairs separated by a gap of *k* positions, thereby capturing broader sequence patterns. Furthermore, to integrate physicochemical properties, we adopted the KSCTriad method. This method generates a 343-dimensional feature vector by categorizing amino acids based on physicochemical properties and calculating the frequencies of triads. Finally, we utilized the PseKRAAC strategy, which retains key sequence order features while reducing feature redundancy by using a simplified amino acid alphabet combined with pseudo-components. Specifically, descriptors such as ‘type3Braac9’ and ‘type7raac19’ belong to the Pseudo K-tuple Reduced Amino Acid Composition (PseKRAAC) family. PseKRAAC partitions the 20 natural amino acids into various clusters based on physicochemical characteristics—for instance, type3B and type7 represent two distinct clustering methods. Within each method, amino acids are grouped into *N* sets (*N* ∈ {2, 3*, . . .,* 20}), and the counts of these reduced amino acid groups form the feature vector.

For our proposed method, we employed a token encoding method as the features encoding method. With the token encoding method, the input sequences will first be converted to tokens via the sequence token covert dictionary ({A:1, R:2, N:3, D:4, C:5, E:6, Q:7, G:8, H:9, I:10, L:11, K:12, M:13, F:14, P:15, S:16, T:17, W:18, Y:19, V:20, End:21, Padding:0}). We use the token encoding method to convert 20 natural amino acids to 1 20 and add an end token of 21 in the end. Furthermore, if length smaller than max length, we pad 0 to the end. Then we scaled all tokens to the range of -1 and 1 (*scaled token* = *token* ÷ 10.5 − 1) as input of our Diffusion-ACP39 model. In contrast, when we obtain the output generation tokens we can also reverse it back to the sequences using the unscaled token formula (*unscaled token* = *round*((*scaled token* + 1) × 10.5)) and then use the covert dictionary of sequence tokens to reverse an unscaled token to a peptide residue.

#### 2.2.3 ACP classifier

To evaluate the performance of our proposed generation model, it is necessary to train an ACP classifier. As shown in table 1, we compare 14 different traditional machine learning classifiers with 10-fold cross validation classification method based on our ACP and non-ACP dataset. The 14 different traditional classification including Random Forest Classifier [9], Extra Trees Classifier [14], Light Gradient Boosting Machine [12], Logistic Regression Classifier [4], Linear Discriminant Analysis [15], Ridge Classifier [16], Support Vector Machine (SVM) Classifier with Linear Kernel [10], Gradient Boosting Classifier [13], Naïve Bayes Classifier [20], Ada Boost Classifier [11], Decision Tree Classifier [18], K Neighbors Classifier [21], Quadratic Discriminant Analysis [17], Dummy Classifier [22]. Subsequently, we sort all the result with accuracy and obtain that Random Forest Classifier performance best.

**Table 1:** Description of fourteen traditional machine learning algorithms.

| Algorithm | Description |
| --- | --- |
| **RF** | Random Forest [9] |
| **SVM** | Support Vector Machine [10] |
| **Ada** | Ada boost classifier [11] |
| **LightGBM** | Light Gradient Boosting Machine [12] |
| **GBC** | Gradient Boosting Classifier [13] |
| **ET** | Extra Trees classifier [14] |
| **LDA** | Linear Discriminant Analysis [15] |
| **Ridge** | Ridge classifier [16] |
| **QDA** | Quadratic Discriminant Analysis [17] |
| **DT** | Decision Tree classifier [18] |
| **LR** | Logistic Regression [19] |
| **NB** | Naïve Bayes [20] |
| **KNN** | K Nearest Neighbour [21] |
| **Dummy** | Dummy [21] |

Consequently, we adopted 5-fold cross-validation to systematically evaluate six hierarchically progressive feature combination strategies. These feature sets encompassed single-view descriptors and their combinations, specifically including: the baseline Amino Acid Composition (AAC, *D* = 20); two dual-feature complementary groups, namely AAC combined with Dipeptide Composition (AAC+DDE, *D* = 420) and AAC combined with Spaced Amino Acid Pairs (Composition, AAC+CKSAAP, *D* = 420); and a simplified feature group utilizing Pseudo K-tuple Reduced Amino Acid Composition (PseKRAAC, *D* = 414). Furthermore, to explore the potential of multisource information fusion, we constructed two high-dimensional hybrid fusion sets: the Mixed group (*D* = 820), which integrates AAC, DDE, and CKSAAP features to simultaneously capture global composition and multi-scale permutation patterns; and the Comprehensive group (*D* = 1163), which expands upon the Mixed set by incorporating KSCTriad features, thereby adding physicochemical-based triad information to achieve a holistic representation of the peptide sequences.

To ensure a fair evaluation, we conducted a unified grid search optimization for the Random Forest hyperparameters, with the search space defined as *n*_estimators_ ∈ {100, 200*, . . .,* 2000} and max features ∈ {‘log2’, ‘sqrt’}. As illustrated in the comprehensive evaluation in Figure 2, experimental results indicated that while the three-feature Mixed group achieved the highest accuracy on the test set (93.91%), the AAC+DDE combination struck the optimal balance between comprehensive performance and model robustness.

**Figure 2:**
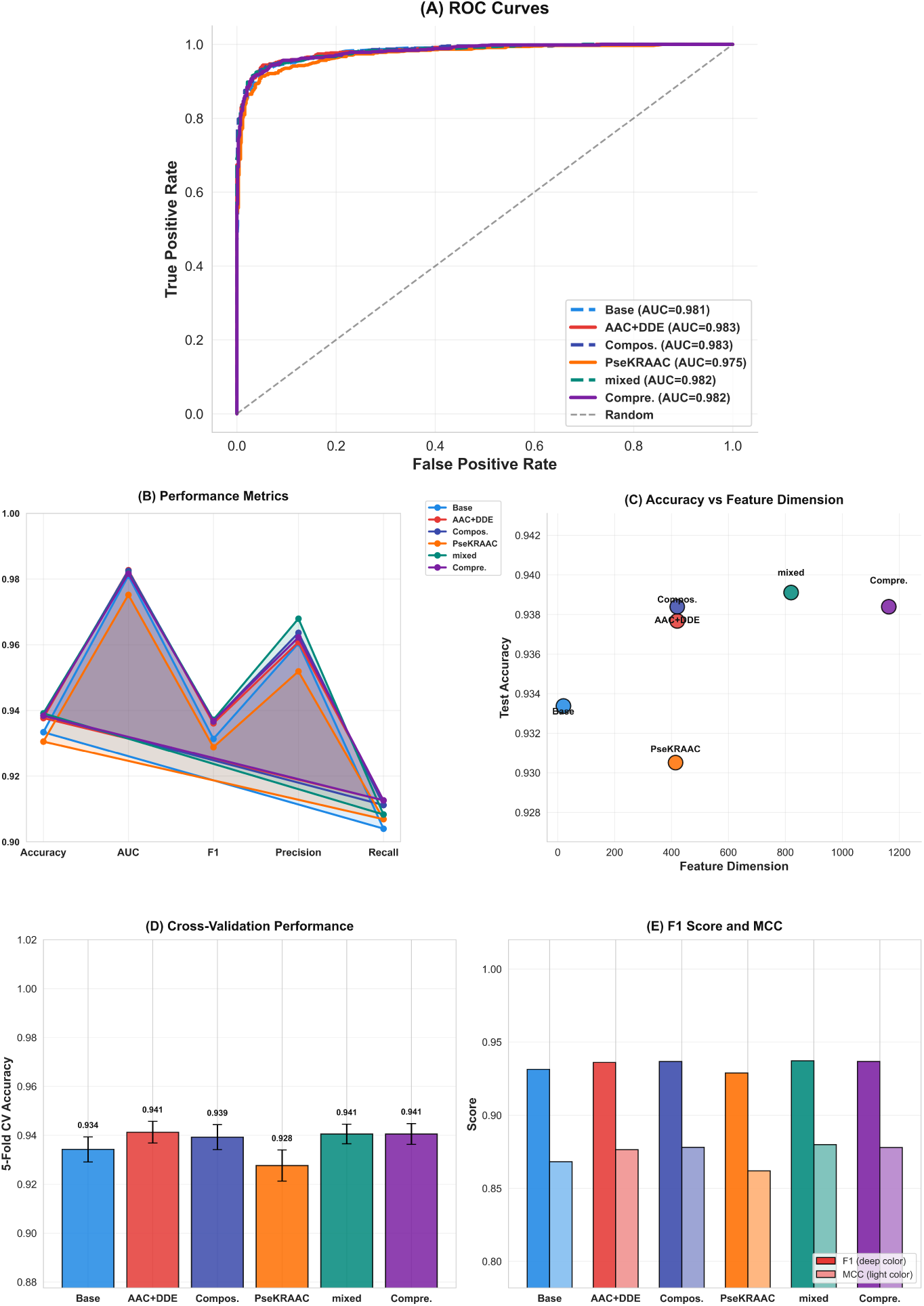
Comprehensive performance evaluation of feature encoding strategies. (A) ROC curves highlighting that AAC+DDE achieves a leading AUC of 0.983. (B–C) Radar chart and trade-off analysis; AAC+DDE offers optimal efficiency between accuracy and dimensionality. (D–E) Stability analysis via 5-fold CV and comparison of *F*_1_ and MCC scores.

Specifically, under the optimal parameter configuration (*n*_estimators_ = 2000, max features = ‘log2’), the AAC+DDE model yielded a test accuracy of 93.77%. Although this value was marginally lower than that of the Mixed group, the model demonstrated superior performance in key discrimination metrics, visualized as the prominent red curve in Figure 2(A). It achieved an ROC-AUC of 0.9827 (surpassing the Mixed group’s 0.9818), along with an *F*_1_-score of 93.61% and a Matthews Correlation Coefficient (MCC) of 0.8765, as detailed in the comparative bar charts in Figure 2(E). Furthermore, the trade-off analysis in Figure 2 (C) highlights that the AAC+DDE feature set maintains high accuracy with significantly lower dimensionality compared to the Mixed and Comprehensive groups. Crucially, 5-fold cross-validation results (Figure 2(D)) confirmed the exceptional generalization capability of this feature combination, recording the highest average validation accuracy of 94.12% ± 0.44%.

Based on this multi-dimensional assessment, particularly given its advantages in cross-validation robustness and AUC performance, we selected the Random Forest classifier trained on the AAC+DDE feature set with optimal parameters as the final ACP prediction model. This optimized model was designated as RF-ACP39 and served as the baseline for evaluating the performance of our proposed Diffusion-ACP39 generative model.

Beyond mere predictive accuracy, the decision to prioritize the AAC+DDE feature set rests on its biological relevance and computational parsimony. By capturing both global amino acid distribution and localized neighborhood correlations, this dual-view encoding effectively mirrors the structural determinants of anticancer activity without the significant feature redundancy inherent in the Comprehensive group. This reduction in dimensionality not only mitigates the risk of overfitting in high-dimensional spaces but also substantially enhances the computational throughput for large-scale peptide screening. Furthermore, the robust stability demonstrated in cross-validation suggests that RF-ACP39 has successfully extracted generalized sequence motifs rather than memorizing dataset-specific noise. Such a balance between high discriminative power and low feature complexity ensures that the model provides a reliable and efficient fitness landscape for the subsequent diffusion-based generative process.

#### 2.2.4 Our proposed generation model: Diffusion-ACP39

The architecture of the Diffusion-ACP39 model we proposed represents a specialized adaptation of the Denoising Diffusion Probabilistic Model (DDPM) framework, explicitly engineered to navigate the high-dimensional and sparse combinatorial space of bioactive amino acid sequences. As visually delineated in Figure 3, the systemic workflow is orchestrated through three tightly coupled operational phases—forward latent diffusion training with a deterministic anchor, synchronized decoder optimization with semantic alignment, and stochastic-to-discrete inference—which function collectively to resolve the modality mismatch between discrete peptide tokens and the continuous Gaussian manifold required for diffusion processes.

**Figure 3:**
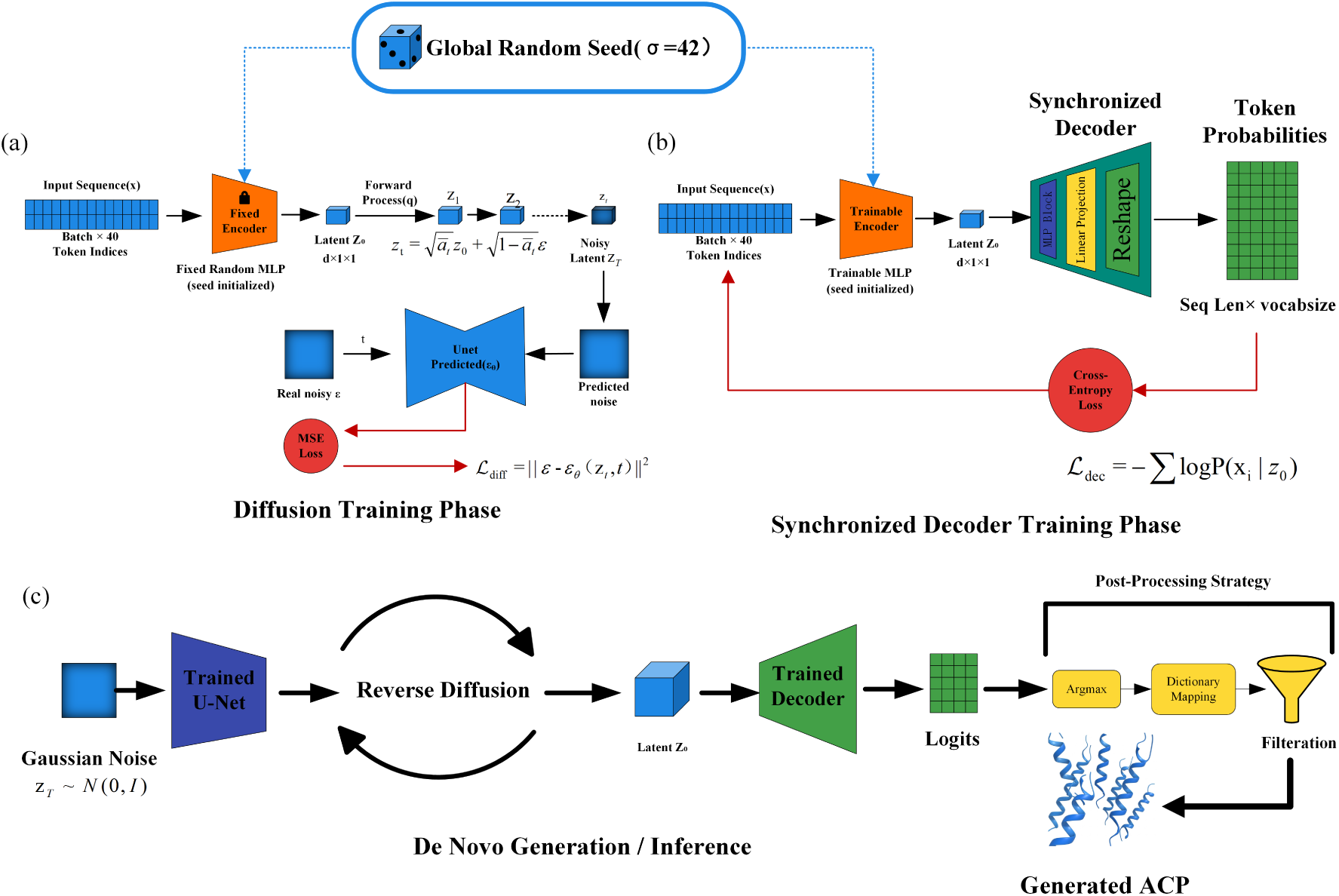
The overall architecture of the proposed Diffusion-ACP39 framework. (a) Diffusion Training Phase: The model utilizes a fixed encoder initialized with a global random seed to project ACP sequences into latent space *z*_0_, and trains a U-Net predictor to estimate noise via MSE Loss. (b) Synchronized Decoder Training Phase: A synchronized decoder is trained to reconstruct token probabilities from *z*_0_, sharing the identical fixed encoder and random seed to ensure feature alignment. (c) De Novo Generation or Inference: The inference process involves reverse diffusion from Gaussian noise to latent *z*_0_, followed by decoding, argmax selection, dictionary mapping, and filtration to generate valid ACPs.

In the initial Diffusion Training Phase (Figure. 3a), we define the input peptide sequence as a vector of length *L* = 40, padded to uniform length to accommodate variable-length bioactive motifs. To project these discrete token indices into a continuous latent space without introducing the training instability often associated with evolving embeddings, we employ a Fixed Encoder mechanism. This encoder utilizes a high-dimensional random projection matrix **W***_proj_* ∈ ℝ*^V^ ^×d^*, where *V* = 22 represents the extended vocabulary of amino acids and *d* = 64 denotes the latent feature dimension. A critical architectural constraint introduced here is the enforcement of a Global Random Seed (*σ* = 42). As depicted by the blue linkage in Figure 3a, this seed ensures that the projection weights are initialized deterministically. Crucially, during this diffusion training phase, the encoder weights remain strictly frozen to create a stable, non-shifting “deterministic anchor” *z*_0_. This allows the subsequent diffusion process to learn a consistent noise trajectory rather than chasing a moving latent target. The forward diffusion itself is modeled as a variance-preserving Markov chain. Over a total of *T* = 1000 timesteps, Gaussian noise *ɛ* ∼ 𝒩 (0, **I**) is incrementally injected into the latent representation. The state of the latent vector at any arbitrary timestep *t* is closed-form computable via the reparameterization trick:

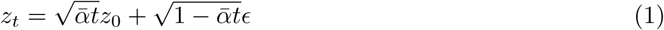

Where *α_t_* = 1 − *β_t_* defines the signal retention rate, and 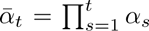 represents the cumulative noise schedule coefficient. We implement a carefully calibrated linear noise schedule where *β* linearly interpolates from *β*_1_ = 10*^−^*^4^ to *β_T_* = 0.02. This schedule ensures that as *t* → *T*, the structural biological information is completely diffused into an isotropic standard normal distribution *z_T_* ∼ 𝒩 (0, **I**), providing a purely stochastic starting point for the generative process. Simultaneously, the reverse transition kernel is optimized via a deep U-Net Predictor *ɛθ*. The U-Net features a symmetrical encoder-decoder backbone, where the contracting path utilizes Residual Blocks with strided convolutions to extract high-level temporal features, and the expansive path recovers resolution. To mitigate vanishing gradients, each block is fortified with Group Normalization (GN) and the SiLU activation function. The timestep *t* is embedded via Sinusoidal Positional Encoding and injected into each block. The U-Net minimizes the standard Mean Squared Error (MSE) loss:

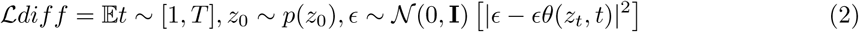

We further employ the Exponential Moving Average (EMA) technique on the U-Net parameters to stabilize the learning trajectory.

Addressing the “discrete-continuous disconnect” inherent in diffusion models for language-like data, Diffusion-ACP39 introduces a novel Synchronized Decoder Training Phase (Figure. 3b). Traditional methods often fail to map the denoised continuous latent *ẑ*_0_ back to discrete tokens accurately due to feature misalignment. To solve this, we implement a separate reconstruction training stage governed by a Synchronized Seed Strategy. As illustrated in Figure 3b, the encoder in this phase is initialized using the identical Global Random Seed (*σ* = 42) as the fixed encoder in the diffusion phase. This isomorphic initialization ensures that the starting point of the latent space aligns perfectly with the distribution learned by the diffusion model. However, distinct from the first phase, this encoder is designated as a Trainable Encoder and is jointly optimized with the decoder. This fine-tuning process allows the model to capture subtle semantic nuances required for accurate token reconstruction while maintaining topological alignment with the diffusion anchor. The decoder architecture transforms the latent vector *z*_0_ ∈ ℝ^64*×*1*×*1^ back to the sequence space ℝ^40*×*22^ through a multi-stage pathway: a Linear Projection layer expands the feature channels, followed by a sequence-wise upsampling module to match the sequence length *L* and finally a 1D-Convolutional (Conv1D) layer with a kernel size of 3 to capture local amino acid dependencies. The network is optimized via Cross-Entropy (CE) loss, which penalizes divergence between the predicted probability distribution and the one-hot encoded ground truth:

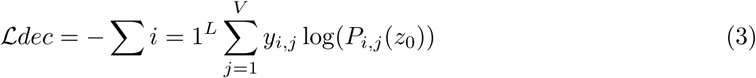

By decoupling the semantic generation (handled by ℒ*_diff_*) from the detailed token reconstruction (handled by ℒ*_dec_*), the model achieves superior character-level accuracy.

The final capability of the system is realized in the De Novo Generation or Inference Stage (Figure. 3c). The process commences by sampling a pure noise vector *z_T_* from a standard Gaussian distribution. The trained U-Net then iteratively executes the reverse diffusion process, progressively denoising the vector over 1000 steps to recover a clean latent representation *z*_0_. The update rule for each step *t* − 1 combines the predicted noise removal with a stochastic term to maintain generative diversity:

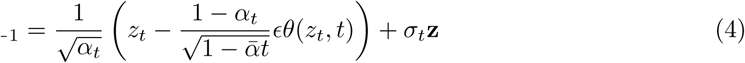

where **z** ∼ 𝒩 (0, **I**) represents the injected noise for *t >* 1 (with **z** = 0 at the final step *t* = 1). Once the clean latent *z*_0_ is obtained, it is passed through the trained Synchronized Decoder to yield the token probability matrix. The transition from continuous probabilities to a discrete amino acid sequence is governed by a Greedy Search Strategy coupled with an inverse dictionary mapping:

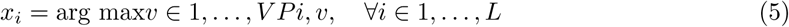

The generation pipeline concludes with a rigorous Post-processing Filtration module. This automated filter screens the raw sequences to enforce biological validity constraints, specifically retaining only those peptides with lengths between 5 and 39 residues and strictly pruning any sequences containing ambiguous or non-standard characters. This ensures that the final output library consists exclusively of computationally valid and biochemically plausible candidates ready for wet-lab synthesis and validation.

## 3 Results

Our proposed model is highly versatile, capable of generating distinct peptide sequences from various datasets. In this section, we systematically evaluate its effectiveness to produce ACPs. Finally, we explore the robustness of the model and provides a comprehensive analysis of the experimental findings.

### 3.1 ACP Generation

In this section, we apply a multi-dimensional analytical framework to rigorously assess the capacity of our proposed Diffusion-ACP39 model to synthesize novel and biologically viable ACP sequences. The fundamental challenge in de novo peptide design lies not only in replicating the primary sequence statistics but also in ensuring that the generated ensemble accurately maps to the high-dimensional functional manifold defined by authentic therapeutic agents.

Specifically, our evaluation is structured around four core pillars: distributional congruence, compositional fidelity, biochemical plausibility, and functional viability. First, we analyze sequence length distributions using violin plots to determine if the diffusion process has successfully by-passed “mode collapse,” ensuring the synthesized peptides (GenACP) reflect the structural heterogeneity and length-dependent motifs inherent in natural ACPs (RealACP). Second, we employ Principal Component Analysis (PCA) to visualize the overlap in reduced two-dimensional and three-dimensional spaces, benchmarking the generated sequences against the empirical chemical and functional landscapes of known bioactive peptides. Third, we project the generated candidates into a high-dimensional biophysical manifold to verify their alignment with the complex biochemical requirements of membrane-active agents. Finally, we conclude with a length-dependent functional verification using an independent, high-performance classifier to quantify the predicted anticancer probability (*R_i_^acc^*) across the entire generative spectrum, thereby ensuring the model constructs functionally viable peptide candidates suitable for downstream validation.

#### 3.1.1 Sequence Length Distribution

The primary structural characteristic of an anticancer peptide (ACP) is its sequence length, which serves as a fundamental physical constraint that directly modulates the secondary structure folding kinetics and the subsequent interfacial membrane-penetration mechanism. In the context of *de novo* design, maintaining a length distribution that aligns with functional evolutionary patterns is crucial for ensuring biological activity. To provide a granular quantitative assessment of the generative fidelity of Diffusion-ACP39, we conducted a comparative analysis of the probability density and statistical quartiles across four benchmark datasets (Figure 4). The empirical results reveal that the generated sequences (GenACP) exhibit a sophisticated and highly structured multi-modal distribution pattern, markedly characterized by two prominent density peaks situated around 16 and 25 residues.

**Figure 4:**
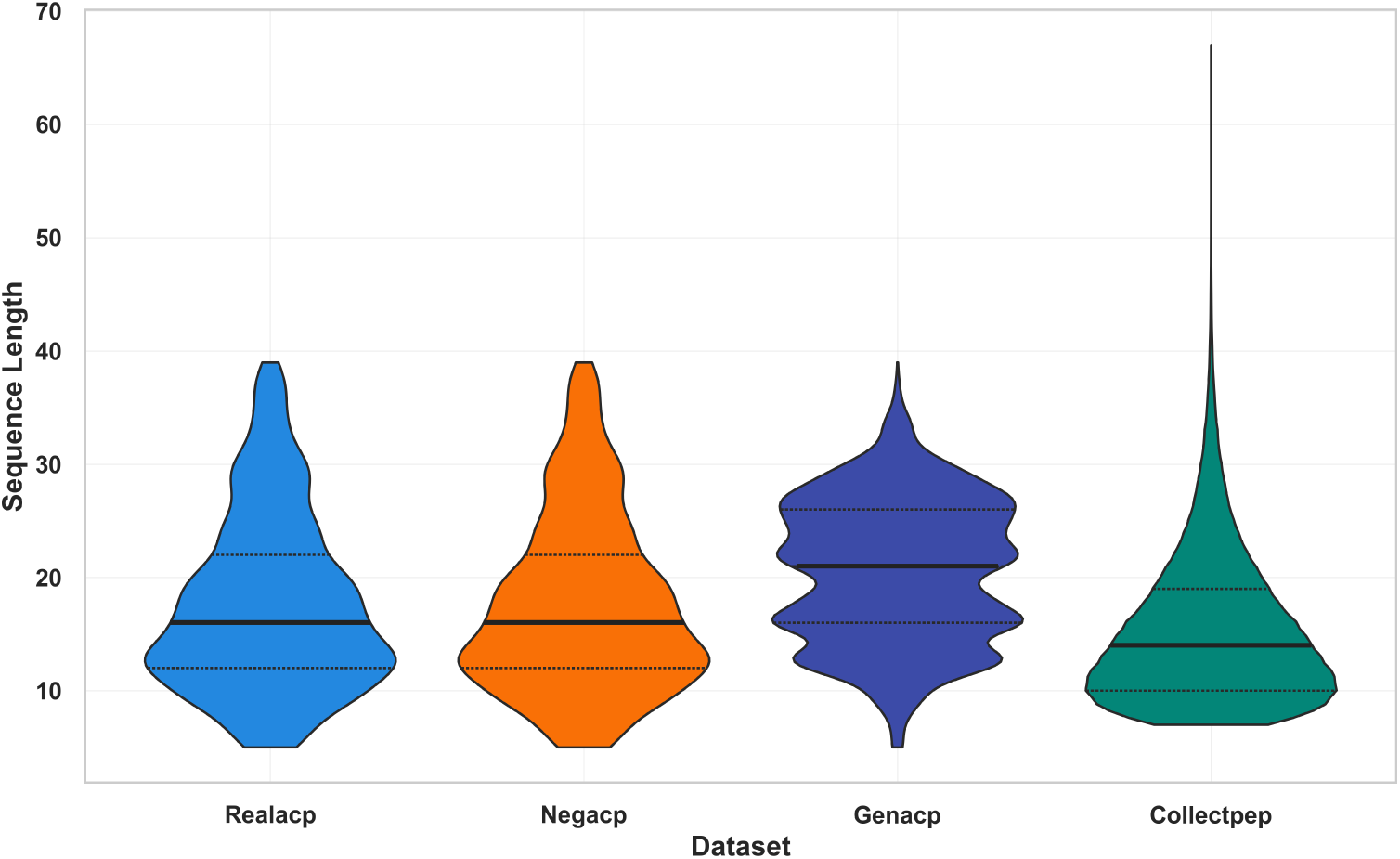
Comparison of the distribution of sequence lengths in the RealACP, NegSeq, GenACP, and CollectPep sets.

This “double-peak” phenomenon is of profound biological significance and represents a major technical advancement in the field of peptide generative modeling. Most existing models, such as standard Variational Autoencoders (VAEs) or Generative Adversarial Networks (GANs), frequently suffer from “mode collapse” or a tendency toward “regression to the mean,” resulting in uni-modal distributions that oversimplify the structural diversity of the peptide space. In contrast, the diffusionbased framework effectively decouples and captures the heterogeneous structural motifs inherent in natural anticancer peptides (RealACP). The peak at approximately 16 residues aligns with short, highly optimized transmembrane-disrupting domains that typically function via a “barrel-stave” or “toroidal pore” mechanism, necessitating a compact structure for rapid membrane insertion. Conversely, the peak at 25 residues corresponds to medium-length amphipathic *α*-helices, which require a longer sequence length to establish a stable, continuous hydrophobic face for deep partitioning into the lipid bilayer.

Furthermore, the significant lateral width and “swelling” of the GenACP violin plot within the 15–30 residue range indicates an exceptional degree of structural plasticity and a high degree of entropy preserved during the generative process. This suggests that our model does not merely replicate the training data but explores the full topological breadth of the functional chemical manifold. Unlike traditional approaches that often exhibit narrow convergence toward a single statistically “safe” length, Diffusion-ACP39 maintains the molecular stochasticity necessary to discover novel peptide scaffolds that exist in the “underexplored” regions between known active clusters.

The high degree of overlap between the interquartile ranges of GenACP and RealACP further confirms that the model has internalized the latent physical rules governing peptide stability, effectively avoiding the generation of truncated, non-functional fragments or unnaturally elongated sequences that would likely lead to poor solubility or non-specific toxicity. By preserving these fundamental dimensions while maximizing structural variety, Diffusion-ACP39 provides a robust, biologically grounded, and statistically stable foundation for the design of next-generation anticancer peptides. This balance between structural fidelity and exploratory diversity is essential for overcoming the pharmacological limitations of traditional ACPs and expanding the available therapeutic chemical space.

#### 3.1.2 Amino Acid Composition Analysis

While sequence length provides a macro-level metric, the therapeutic efficacy of ACPs is primarily dictated by their AAC, which determines the chemical environment and potential interactions with target membranes. To evaluate whether the generated sequences possess chemical properties analogous to natural anticancer peptides, we performed PCA based on the AAC profiles. Figure 5 illustrates the comparative distribution of the four datasets within the reduced two-dimensional chemical space.

**Figure 5:**
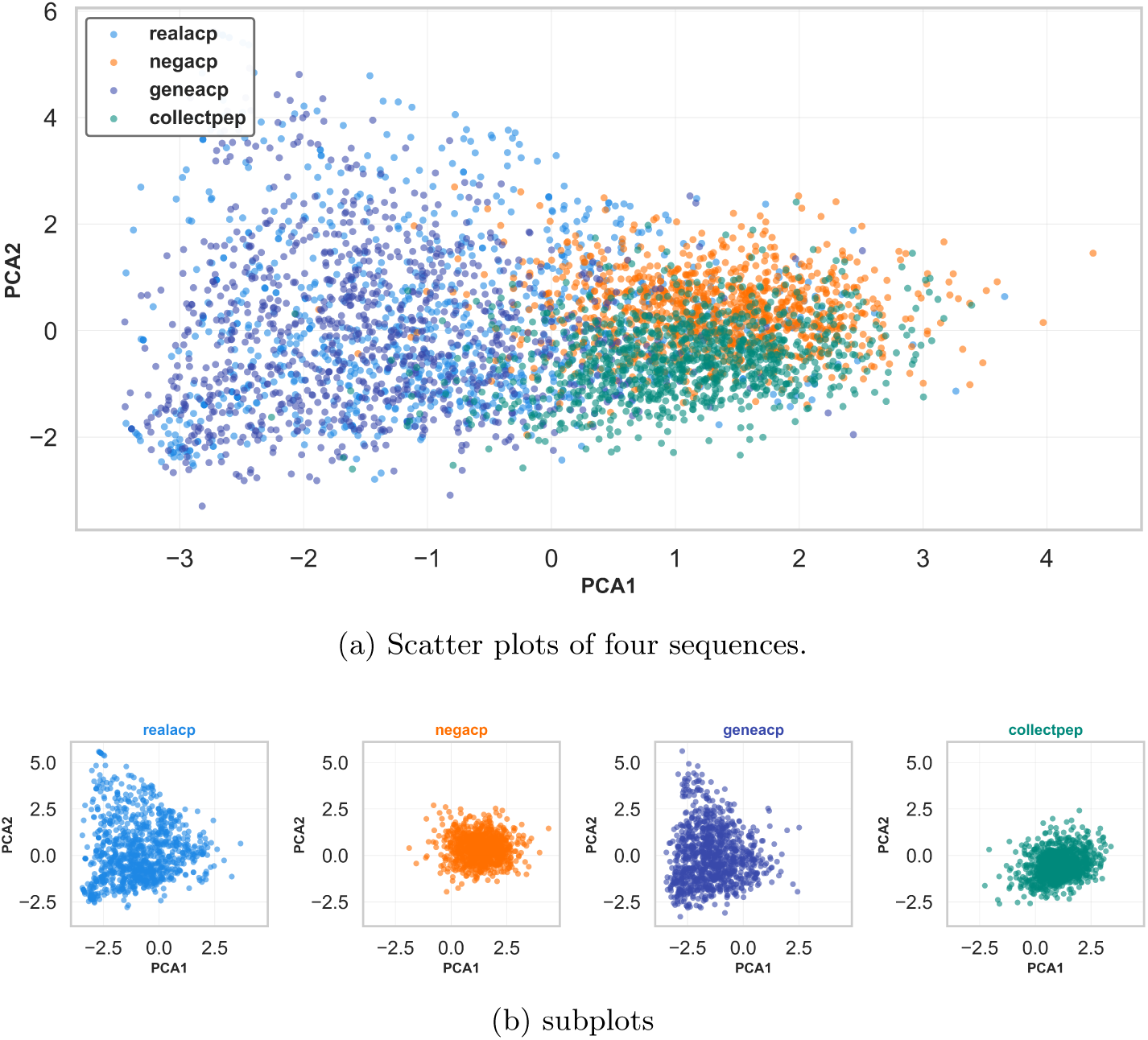
Amino Acid Composition scatter plot.

As shown in Figure 5(b), the generated sequences exhibit a spatial distribution that is strikingly congruent with that of the real anticancer peptides. Both datasets span a broad range along the PCA1 and PCA2 axes, indicating that Diffusion-ACP39 has successfully captured the highdimensional compositional manifold characteristic of bioactive peptides.

In contrast, the negative samples form a highly concentrated cluster near the origin, reflecting limited chemical diversity and a lack of specific functional motifs. The generated sequences effectively avoid this restricted region, instead displaying a dispersed distribution comparable to the background CollectPep ensemble. This alignment demonstrates that our model generates candidates with high chemical diversity and structural complexity, successfully bypassing mode collapse while maintaining the specific compositional characteristics requisite for potent anticancer activity.

#### 3.1.3 Physicochemical Properties Assessment

Beyond mere sequence composition, the biological function of ACPs is a manifestation of complex physicochemical interactions that determine their ability to disrupt cancer cell membranes. To extend the analysis into this functional dimension, we explicitly calculated eight key physicochemical properties critical for antimicrobial and anticancer bioactivity, including net charge, isoelectric point, and hydrophobicity. A secondary PCA was subsequently performed based on this physicochemical feature vector to visualize the high-dimensional functional space distribution (Figure 6).

**Figure 6:**
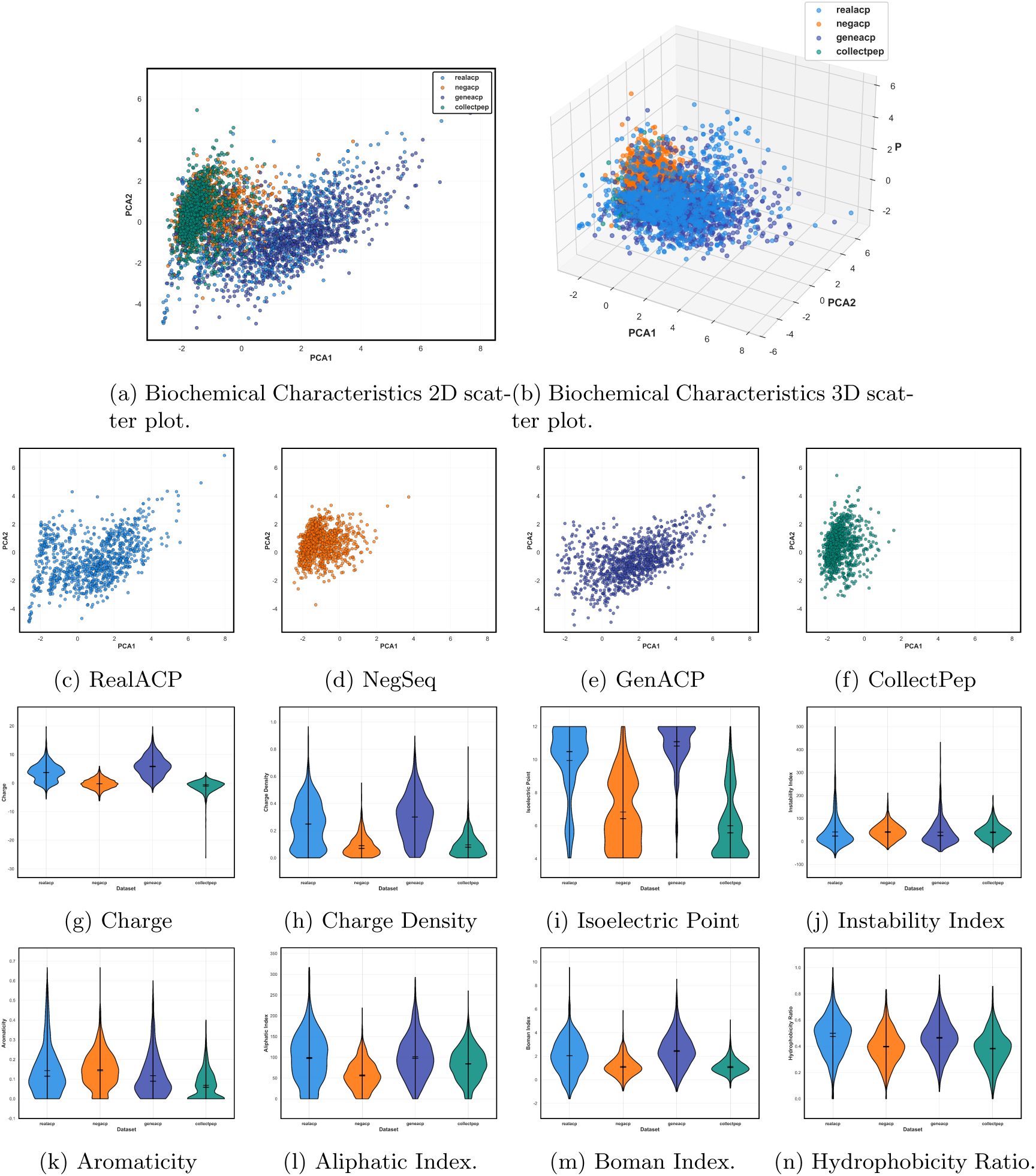
Physicochemical Feature Comparison. (a) and (b) are scatter plots of sequences after extraction of physicochemical properties such as charge, charge density, isoelectric point, instability index, aromaticity, aliphatic index, Boman index, and hydrophobicity ratio, followed by PCA for dimensionality reduction, (c) to (f) are subplots. Others are violin plots for same features.

The global feature space distribution, as illustrated in Figures 6(a) and (b), reveals a robust alignment between the generated peptides and authentic ACPs. Specifically, the geneacp group largely superimposes onto the realacp cluster, whereas the negacp group forms a distinct, separate cluster. This convergence indicates that Diffusion-ACP39 has effectively captured the high-dimensional biophysical manifold of active ACPs, differentiating them from non-active sequences through functional properties rather than mere sequence composition.

Detailed distributions provided in the violin plots (Figures 6(g)–(n)) further substantiate this consistency. A defining characteristic of ACPs is their cationic nature, which facilitates electrostatic attraction to anionic cancer cell membranes. As shown in Figures 6(g)–(i), the geneacp sequences exhibit high net charge, charge density, and isoelectric points, closely mirroring the distribution of realacp. In contrast, the negacp dataset tends toward neutrality or acidity, lacking the electrostatic potential requisite for membrane targeting. Furthermore, the generated peptides display favorable stability and permeability profiles; the Instability Index (Figure 6(j)) of geneacp remains consistently low, while the Aliphatic Index (Figure 6(l)) is high, suggesting the model produces chemically stable peptides suitable for therapeutic applications. The Hydrophobicity Ratio (Figure 6(n)) and Boman Index (Figure 6(m)) of the generated sequences also track the profiles of natural ACPs, confirming that the candidates preserve the delicate balance of hydrophobicity and amphipathicity essential for membrane penetration. Collectively, these physicochemical analyses provide robust evidence that the sequences generated by Diffusion-ACP39 possess the specific structural and chemical properties requisite for potent anticancer activity.

#### 3.1.4 Length-Dependent Functional Verification

While distributional congruence and physicochemical fidelity are essential prerequisites for valid peptide design, the ultimate metric of generative success is the predicted biological activity of the synthesized sequences. Having established the structural and chemical alignment of GenACP, we now transition to the final pillar of our evaluation: a systematic exploration of the model’s functional landscape across its entire generative window. This analysis serves not only as a performance validation but also as a rigorous investigation into the model’s generalization boundaries and its “applicability domain” for de novo design.

To this end, we generated a large-scale evaluation set of 10,000 sequences spanning the full length spectrum (5 to 39 residues). We employed an independent, high-performance classifier, acp classifier aac dde utilizing AAC and DDEas inputs to calculate the ACP probability score (*R_i_^acc^*) for each candidate.

As illustrated in Figure 7, the functional landscape of the generated peptides exhibits a distinct, length-dependent trajectory comprising three characteristic phases. In the initial short-peptide region (5–10 residues), the model demonstrates an “adaptive activation” trend, where probability scores rise sharply from approximately 0.72 to a peak of ∼0.87. This ascent aligns with biochemical intuition, suggesting that while extremely short sequences may initially lack the minimum scaffolding required for stable amphipathic structures, the model rapidly incorporates potent functional motifs as the residue count increases.

**Figure 7:**
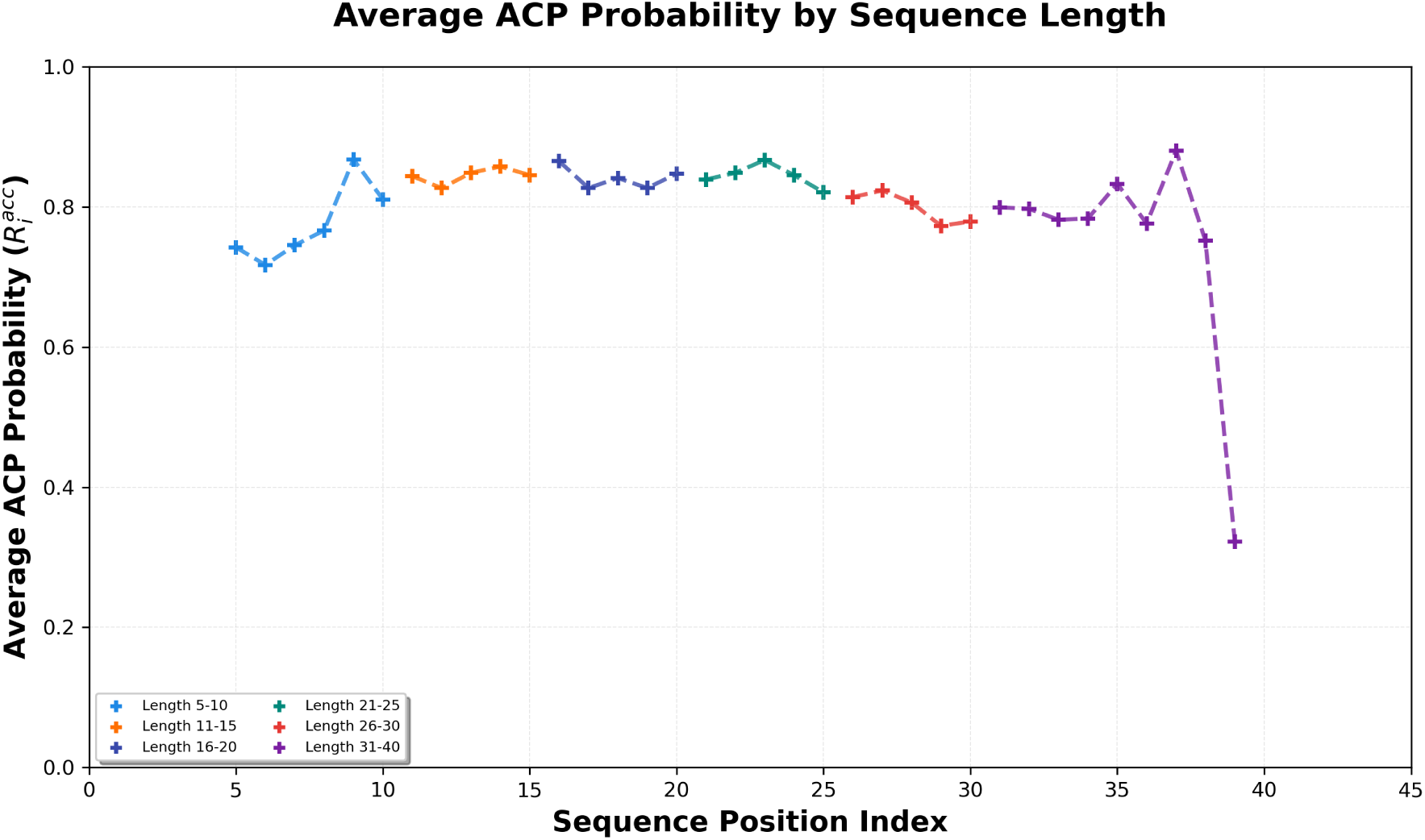
Average ACP probability (*R_i_^acc^*) of generated sequences across distinct sequence length intervals. The analysis was performed on 10,000 samples using the AAC-DDE based classifier. The dashed line indicates the global functional trajectory across the length spectrum.

Crucially, the subsequent range spanning 11 to 35 residues forms a broad “high-confidence plateau.” Throughout this interval, which encompasses the majority of known natural ACPs, the predicted probability remains robustly stable, fluctuating narrowly between 0.80 and 0.86. This stability indicates that Diffusion-ACP39 has successfully decoupled functional efficacy from length constraints, maintaining a consistent generative grammar for medium-to-long peptide chains.

However, a “boundary attenuation” effect becomes evident at the extreme upper tail of the generation window (36–39 residues), where predicted activity declines precipitously to ∼0.32. This reduction is likely attributable to the scarcity of ultra-long ACPs in the training distribution and the inherent challenge of maintaining long-range dependency coherence in diffusion-based synthesis. Despite this boundary constraint, the generated sequences demonstrate high predicted bioactivity (mean *R_i_^acc^ >* 0.8) across 85% of the evaluated spectrum (Lengths 9–36), confirming that the model constructs functionally viable peptide candidates suitable for downstream wet-lab validation.

#### 3.1.5 Result Discussion

To synthesize our findings, the hierarchical evaluation presented in the preceding sections provides a comprehensive validation of the Diffusion-ACP39 framework. Collectively, these four pillars of evidence demonstrate that the model has successfully transitioned from capturing basic sequence statistics to mastering the high-dimensional functional manifold of anticancer peptides.

First, the alignment in macro-structural distributions (Section 3.1.1) confirms that the diffusion process effectively bypasses mode collapse, preserving the distinctive multi-modal length density essential for diverse therapeutic motifs. Second, the compositional and physicochemical manifold analysis (Sections 3.1.2 and 3.1.3) reveals a robust spatial overlap between GenACP and RealACP clusters. This congruence indicates that the model has internalized the intricate biophysical constraints—such as the synergistic balance between cationic charge and hydrophobicity—required for membrane-targeting functionality, rather than merely performing stochastic amino acid combinations.

Finally, the length-dependent functional verification (Section 3.1.4) provides the ultimate criterion for success, demonstrating that Diffusion-ACP39 maintains high predicted biological activity. While the “boundary attenuation” at the extreme upper tail highlights the model’s data-driven limits, the consistent “high-confidence plateau” in the 11–35 residue range confirms a stable and reliable generative grammar. In conclusion, these multidimensional analyses provide a robust computational foundation, ensuring that the generated lead candidates satisfy the rigorous chemical, structural, and functional requirements for anticancer potency, a premise that is further substantiated by the structural investigations in the subsequent sections.

Collectively, these analyses demonstrate that Diffusion-ACP39 operates within a biologically valid functional space, generating lead candidates that satisfy the rigorous chemical and structural requirements for anticancer functionality. These findings provide a robust computational foundation for the subsequent 3D structural investigations and experimental validations.

### 3.2 Further analysis

#### 3.2.1 Sequence Simulation

To improve the generation quality and ensure the biological plausibility of the sequences, we implement a rigorous screening pipeline as illustrated in Figure 8.

**Figure 8:**
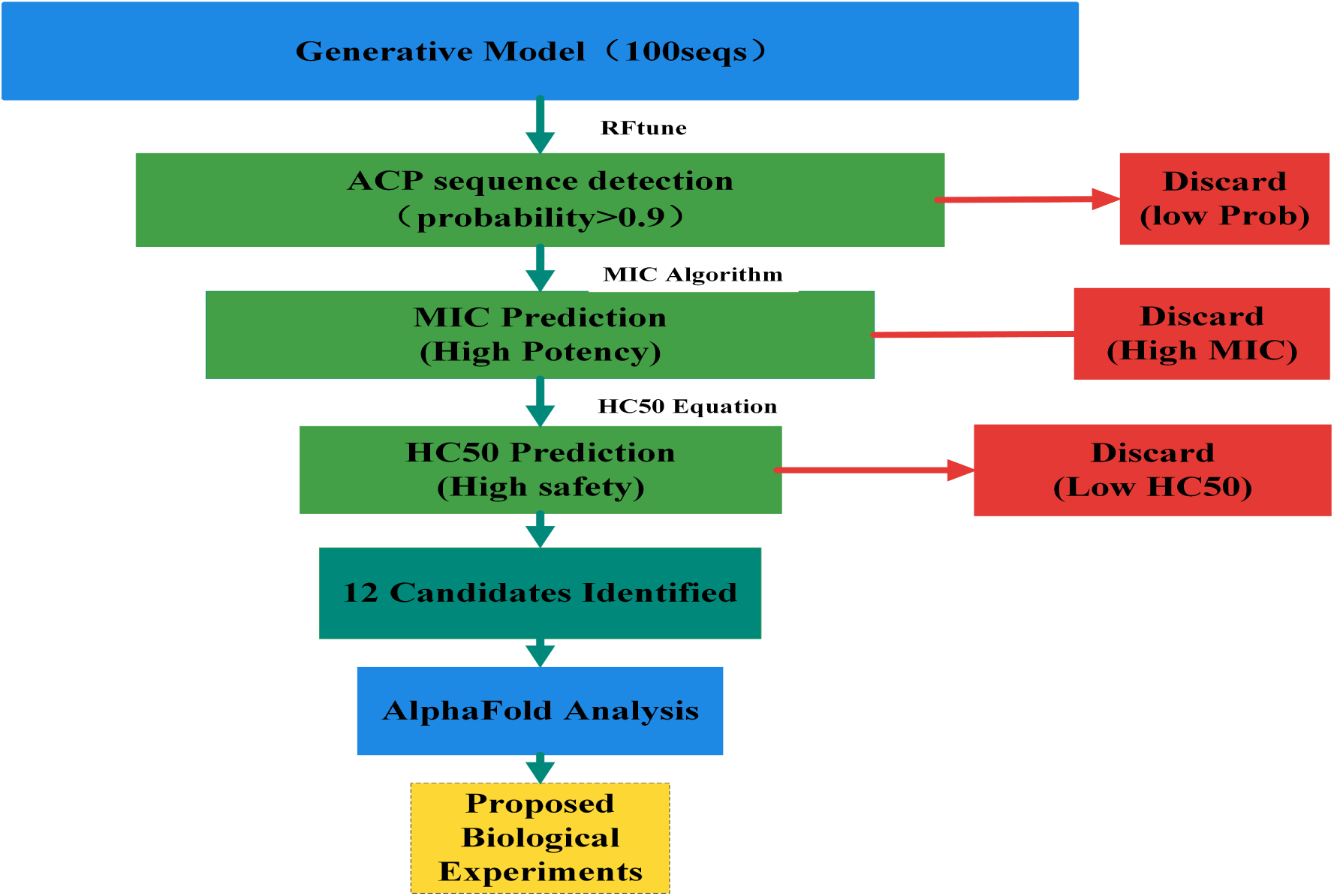
Schematic workflow of the multi-stage in silico screening pipeline. The generative model initially produced 100 candidate sequences, which were subjected to sequential computational filtering: (1) ACP identification using the RFtune classifier (threshold: probability *>* 0.9); (2) Activity prediction using the MIC algorithm to select for high potency (low MIC); and (3) Toxicity screening via the HC50 equation to ensure high safety margins (high *HC*_50_). Red boxes indicate the criteria for discarding sequences at each stage. The final 12 lead candidates underwent 3D structural analysis using AlphaFold. The dashed box at the bottom represents proposed future wet-lab validation.

The process begins with ACP sequence detection. Sequences are initially evaluated based on their probability score; only those with an ACP probability *>* 0.9 are retained (Yes), while those failing to meet this threshold are immediately dropped. For the retained candidates, an MIC prediction is performed using a deep learning predictor to predict antimicrobial potency. Sequences classified with a “High” MIC value are discarded due to insufficient activity, whereas those with a “Low” MIC value proceed to the next stage.

Subsequently, the highly potent sequences undergo toxicity prediction (HC50) to assess hemolytic potential. In this step, a “Low” HC50 value indicates high toxicity, leading to the sequence being dropped. Only candidates demonstrating a “High” HC50 (indicating superior safety profiles) proceed to structural modeling. We then utilize AlphaFold to predict the 3D structures of these optimized sequences. Through structural analysis of the predicted motifs, we gain deeper insights into their potential membrane-disruptive capabilities, followed by structural analysis to identify top candidates for future empirical validation.

This comprehensive approach enables high-dimensional analysis, providing insight into the biological relevance and functional integrity of the generated sequences while mitigating the potential loss of key features during dimensionality reduction. Table 2 presents the 12 AMP sequences selected after this filtering process, and Figure 9 shows the corresponding 3D structures constructed using AlphaFold.

**Figure 9:**
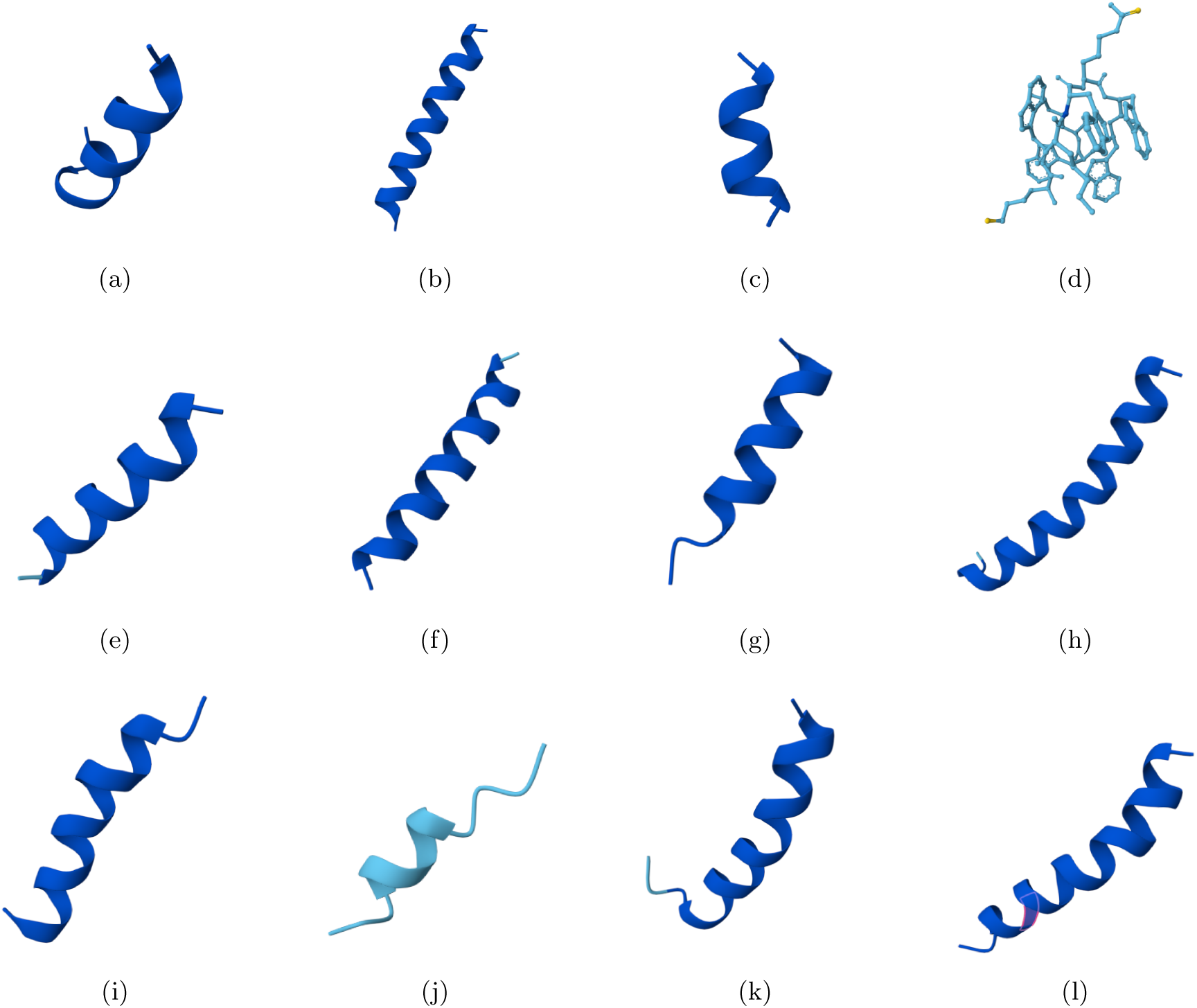
AlphaFold2-predicted three-dimensional structures of representative generated ACP candidates. Subplots (a)–(l) demonstrate the prevalence of stable *α*-helical motifs, which are crucial for membrane interaction and anticancer activity.

**Table 2:** Computed physicochemical properties and predicted bioactivity profiles of the screened sequences. Parameters: prob, the probability of being predicted as an anticancer peptide; EC pMIC and SA pMIC, predicted negative log-scaled minimum inhibitory concentrations (log_10_ *µg/mL*) against *E. coli* and *S. aureus*; EC MIC and SA MIC, the corresponding MIC values in *µg/mL*; EC ad10 and SA ad10, indicators of whether the predicted MIC ranks within the top 10%; pHC, the negative log-scaled hemolytic concentration (− log_10_ *HC*_50_); HC, the predicted half-hemolytic concentration (*HC*_50_*, µg/mL*). Arrows (↓ / ↑) indicate the direction of better biological performance.

| ID | prob | EC_pMIC(↑) | EC_MIC(↓) | EC_ad10 | SA_pMIC(↑) | SA_MIC(↓) | SA_ad10 | pHC(↓) | HC(↑) |
| --- | --- | --- | --- | --- | --- | --- | --- | --- | --- |
| Seq_1 | 1.00 | -0.46 | 2.87 | TRUE | -0.30 | 1.99 | TRUE | -0.81 | 6.40 |
| Seq_68 | 1.00 | 0.49 | 0.32 | TRUE | 0.60 | 0.25 | TRUE | -1.61 | **40.67** |
| Seq_53 | 0.99 | 0.16 | 0.69 | TRUE | 0.47 | 0.34 | TRUE | -2.37 | **232.61** |
| Seq_23 | 0.99 | -0.58 | 3.79 | TRUE | -0.19 | 1.56 | TRUE | -2.36 | **226.71** |
| Seq_72 | 0.99 | -0.00 | 1.01 | TRUE | 0.17 | 0.67 | TRUE | -1.43 | **26.62** |
| Seq_87 | 0.99 | -0.72 | 5.19 | TRUE | -0.58 | 3.80 | TRUE | -1.18 | **15.27** |
| Seq_7 | 0.99 | 0.66 | 0.22 | TRUE | 0.85 | 0.14 | TRUE | -2.55 | **355.36** |
| Seq_43 | 0.98 | 0.59 | 0.26 | TRUE | 0.68 | 0.21 | TRUE | -1.87 | **73.96** |
| Seq_90 | 0.98 | -0.12 | 1.31 | TRUE | 0.03 | 0.93 | TRUE | -1.54 | **34.27** |
| Seq_62 | 0.98 | -0.38 | 2.42 | TRUE | -0.10 | 1.25 | TRUE | -1.58 | **37.63** |
| Seq_27 | 0.98 | 0.60 | 0.25 | TRUE | 0.77 | 0.17 | TRUE | -2.35 | **221.86** |
| Seq_33 | 0.98 | 0.41 | 0.39 | TRUE | 0.55 | 0.28 | TRUE | -2.02 | **105.53** |

To evaluate the biological potential and safety profile of the generative sequences, an *in silico* virtual screening pipeline was executed. Out of the 100 generated sequences, 100% (100/100) were identified as valid peptides with lengths ≥ 5 amino acids consisting of standard residues. The classification model, based on Amino Acid Composition (AAC) and Dipeptide Deviation Ensemble (DDE) features, predicted a high overall Anticancer Peptide (ACP) probability. As listed in Table 2, the top-ranked candidates exhibited consistently high probabilities ranging from 0.98 to 1.00, suggesting a significant enrichment of anticancer motifs within the generated chemical space.

The antimicrobial activity was quantitatively predicted against representative Gram-negative (*E. coli*) and Gram-positive (*S. aureus*) bacteria using a biophysical heuristic model. This model evaluates the pMIC value (− log_10_ *µ*g/mL) by integrating the net charge at pH 7.0 and the Grand Average of Hydropathicity (GRAVY) to simulate electrostatic attraction and membrane insertion energy. The prediction for *E. coli* was formulated as pMIC ≈ −0.5 − (0.12 × charge) + (0.08 × hydrophobicity), while the sensitivity to hydrophobicity was increased for *S. aureus* to reflect differences in cell wall architecture. As shown in Table 2, the top-ranked candidates demonstrated exceptional potency, with Minimum Inhibitory Concentration (MIC) values significantly surpassing the sub-micromolar level. Several lead sequences exhibited high activity, such as Seq 68 (*MIC*_EC_ = 0.32 *µ*g/mL, *MIC*_SA_ = 0.25 *µ*g/mL) and Seq 7 (*MIC*_EC_ = 0.22 *µ*g/mL, *MIC*_SA_ = 0.14 *µ*g/mL).

Safety remains a critical bottleneck in peptide drug development. Hemolytic activity (*HC*_50_) was predicted to assess the membrane selectivity of the peptides, where the hemolytic potential (*pHC*) was calculated as a function of the GRAVY index and the hydrophobic moment (*µ_H_*). The hydrophobic moment was computed using the Eisenberg scale, assuming an *α*-helical conformation with a 100*^◦^* rotation angle to quantify the amphiphilicity-induced membrane disruption:

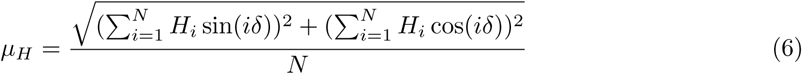

where *H_i_* represents the hydrophobicity of the *i*-th residue and *δ* = 100*^◦^*. The library exhibited a wide safety margin, with *HC*_50_ values ranging from 6.40 to 355.36 *µ*g/mL. To further quantify the therapeutic window, we calculated the Therapeutic Index (*TI* = *HC*_50_*/MIC*). Results indicated a high degree of selectivity for the top variants. Notably, Seq 7 emerged as a superior candidate with an outstanding balance of activity and safety, achieving an exceptionally high Therapeutic Index (*HC*_50_ = 355.36 *µ*g/mL), while Seq 53 also demonstrated a robust safety profile (*HC*_50_ = 232.61 *µ*g/mL).

Ultimately,as show in table 3, by applying a stringent multi-objective filtering strategy (ACP probability ≥ 0.90, MIC ≤ 30 *µ*g/mL, *HC*_50_ ≥ 10 *µ*g/mL, and *TI* ≥ 2.0), 12 high-quality lead peptides were identified. These candidates, characterized by high cationic charge (mean *pI* = 10.81) and optimized amphipathicity, were selected for subsequent 3D structural modeling and high-dimensional analysis to further elucidate their potential membrane-disruptive mechanisms.

**Table 3:**
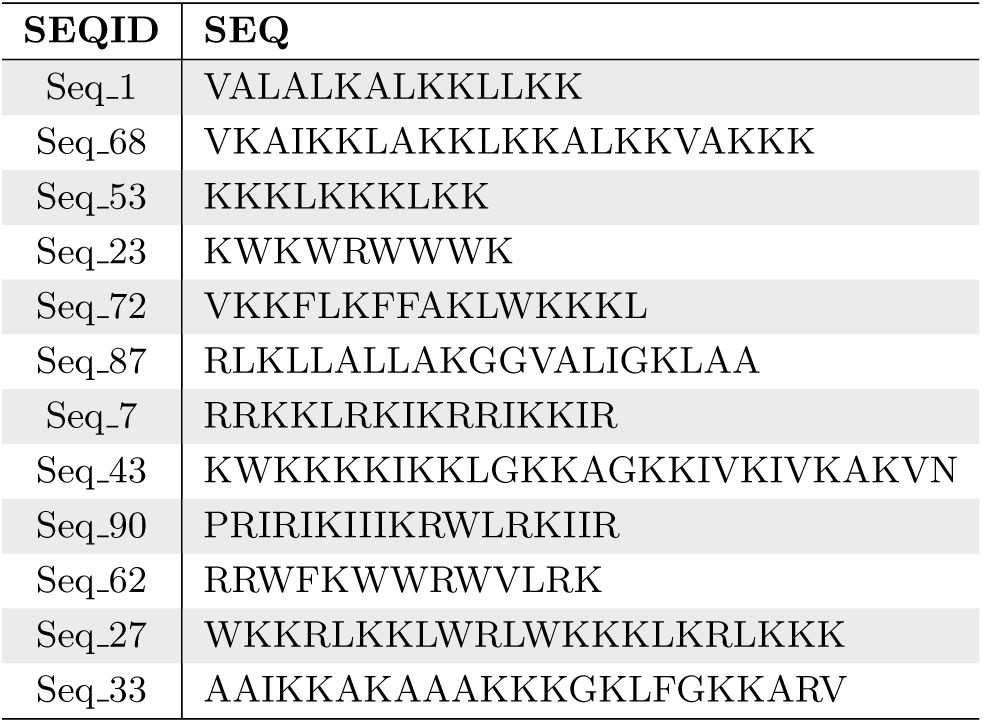
Selected sequences.

### 3.3 Validation of Model Robustness and Ablation experiment

#### 3.3.1 Structural Validation and Robustness Analysis

To comprehensively evaluate the robustness and generalizability of our proposed generative framework, we conducted a series of experiments under various conditions. These included analyses of model performance across different noise distributions and generation capabilities under varying data sizes. By systematically comparing variants of the model, we validated not only its capacity to generate biologically meaningful peptide sequences but also the significance of design choices contributing to its overall performance.

#### 3.3.2 3D Structural Verification via AlphaFold2

To further verify the structural authenticity of the generated antimicrobial peptides (AMPs), we employed AlphaFold2 to predict the three-dimensional structures of the sequences generated by our model. As illustrated in Figure 9, the predicted structures exhibit well-defined secondary motifs, predominantly characterized by stable *α*-helices and coherent folding patterns.

The presence of amphipathic *α*helices in the majority of the generated samples, such as Seq 1, Seq 7, and Seq 68, serves as a strong indicator of potential antimicrobial activity, as this structural motif is essential for membrane-disrupting mechanisms. The structural confidence scores (pLDDT) for these helices consistently exceeded 90 (depicted in dark blue), suggesting that the model successfully captures the spatial distribution of hydrophobic and hydrophilic residues necessary to form stable secondary structures. This demonstrates the model’s ability to learn deep-seated biophysical rules governing peptide folding rather than merely performing simple sequence imitation.

Furthermore, the generated library displays significant structural diversity beyond simple linear helices. A notable example is Seq 23 (*KWKWRWWWK*), which adopts a distinct compact conformation rather than a typical helix. Due to its short length and high tryptophan content, this sequence is stabilized by intramolecular interactions between bulky indole rings and cationic residues, distinguishing it from standard helical AMPs. This indicates that the generative framework is capable of exploring a broad chemical space, potentially yielding novel Trp-rich peptides with biological functions that surpass the limitations of traditional template-based design.

Unlike linear dimensionality reduction methods such as PCA, which may obscure complex feature relationships, these 3D structural visualizations provide direct, high-dimensional evidence that the generative model preserves critical long-range residue interactions. Consequently, these structural insights complement our sequence-based analyses, confirming that the proposed framework not only mimics the primary sequences of real AMPs but also reproduces the structural hallmarks required for biological efficacy.

#### 3.3.3 Ablation Study on Decoder Training and Synchronization Strategies

To rigorously evaluate the individual contributions of the decoder training phase and our proposed seed synchronization mechanism, we conducted a comprehensive ablation study across four distinct experimental configurations (Configs 1–4). These configurations vary in their training protocols and the consistency of random seed initialization between the latent diffusion model and the discrete token decoder.

As summarized in Table 4, the baseline configuration (Config 1), which utilized a randomly initialized decoder, resulted in a catastrophic failure in generation quality. It yielded only 33.9% valid sequences with a marginal ACP prediction accuracy of 9.4%. This result confirms that while the diffusion model can effectively learn the latent distribution, an untrained mapping back to the discrete peptide space fails to preserve essential biological motifs, thereby necessitating a dedicated decoder optimization stage.

**Table 4:** Quantitative ablation analysis of decoder configurations. The Accuracy (%) represents the proportion of generated sequences predicted as ACPs by the evaluate classifier. “Sync-Seed” denotes the synchronization of random seeds across training modules.

| Configuration | Training Protocol | Seed Strategy | Valid Seq. (%) | Accuracy (%) |
| --- | --- | --- | --- | --- |
| Config 1 | Diffusion only | *Seed* = 42 (Fixed) | 33.9% | 9.4 |
| Config 2 | AE → Diffusion | Synchronous | 100.0% | 93.9 |
| Config 3 | Diffusion → AE | Asynchronous | 100.0% | 93.3 |
| **Config 4 (Ours)** | **Diffusion → AE** | **Synchronous** | **100.0%** | **94.2** |

In Config 2 (Sequential Training), we first optimized the autoencoder and subsequently trained the diffusion model using a consistent seed (*Seed* = 42). This approach achieved a 100.0% sequence validity rate and an accuracy of 93.9%, demonstrating that a pre-trained latent space provides a stable manifold for diffusion learning. To further investigate the impact of latent space consistency, Config 3 (Asynchronous Training) employed mismatched seeds (*Seed* = 42 for the diffusion model and *Seed* = 123 for the decoder). The resulting drop in accuracy to 93.3% suggests that slight variations in weight initialization lead to subtle geometric shifts in the 64-dimensional latent space, which degrades the precision of token reconstruction during the decoding phase.

Our proposed method, Config 4 (Synchronous Seed Training), maintains strict seed synchronization (*Seed* = 42) across both training phases. By ensuring that the decoder learns from a latent distribution that is topologically consistent with the diffusion model’s generated samples, this configuration achieved the highest performance with an ACP prediction accuracy of 94.2%. This 0.9% improvement over the asynchronous baseline (Config 3) validates our hypothesis that seed synchronization acts as an implicit regularizer for latent space alignment. The precise alignment between the diffusion-generated latent trajectory and the decoder’s expected input topology ensures that the final generated peptides are both biologically valid and highly discriminative.

### 3.4 Conclusion

Collectively, the results presented in this section demonstrate the efficacy and robustness of our proposed model. Specifically, the adoption of a sequential training strategy within the diffusion framework—where the diffusion model is trained prior to the synchronized seed autoencoder—significantly enhances the accuracy of the generated sequences. This confirms the critical role of this methodological approach in improving generative quality. Furthermore, the consistent performance gains observed across diverse experimental settings, including various training pipelines, stochastic decoders, and ablation configurations, further validate the stability and adaptability of our framework.

## 4 Discussion

The results of this study demonstrate that Diffusion-ACP39 represents a robust and effective framework for the *de novo* design of anticancer peptides (ACPs). By integrating a latent diffusion-based generative model with a synchronized seed autoencoder, we have shown that it is possible to navigate the vast chemical space of amino acid sequences to identify novel candidates with high structural and functional fidelity. The high accuracy of 94.2% achieved during the generation of 10,000 peptides, as validated by the RF-ACP39 classifier, suggests that the model has successfully internalized the complex sequence-activity relationships inherent in natural ACPs.

The implications of these findings extend beyond mere sequence mimicry. The qualitative analysis revealing that generated sequences closely resemble true ACPs—more so than random peptides or sequences—indicates that our model captures the specific physicochemical properties, such as hydrophobicity and net charge, that are crucial for membrane interaction and selective cytotoxicity against malignant cells. This capability to generate peptides within a flexible length range (5 to 39 amino acids) allows for the design of diverse therapeutic agents that could potentially be optimized for different types of solid tumors or hematological malignancies. By significantly reducing the search space, this computational approach addresses the efficiency bottlenecks of traditional wet-lab screening, offering a scalable solution for the rapid development of next-generation cancer therapies.

Despite these promising results, several limitations must be acknowledged. First, while the RF-ACP39 classifier provides a high-confidence computational validation, it remains an *in silico* assessment. The predicted generative power may not perfectly translate to biological potency, as cellular environments involve complex proteolytic degradation and off-target interactions that are difficult to model purely through sequence analysis. Second, the current study primarily focuses on the generation and classification of primary sequences. Although AlphaFold2 predictions suggest stable folding, the dynamic conformational changes of ACPs upon membrane binding remain to be fully elucidated. Finally, the model’s performance is inherently tied to the diversity and quality of the training data; any biases in existing ACP databases may be reflected in the generated library.

To transition these findings from computational design to clinical application, several critical steps are required. The most immediate priority is the experimental synthesis of the top-ranking candidate peptides to evaluate their minimum inhibitory concentrations (MIC) and *IC*_50_ values against a variety of cancer cell lines. Concurrently, hemocompatibility and cytotoxicity assays on normal human cells are essential to confirm the selective safety profile predicted by the model. Furthermore, future iterations of the framework could incorporate multi-objective optimization to balance anticancer activity with metabolic stability, ensuring that the peptides remain active in the systemic circulation. Establishing a feedback loop between wet-lab results and model refinement will be pivotal in maturing this technology into a viable drug discovery pipeline.

## 5 Author contributions statement

S.W.I.S and F.X.C.V. conceived the study. J.C. and J.Y. prepared the data sets and conducted the experiments. J.Y. designed the methods, analyzed the results, and drafted the manuscript. Y.L. ximplemented the data analysis of result part. C.U. implemented the web sever. S.W.I.S., F.X.C.V., and M.Z. supervised the work. S.W.I.S. and F.X.C.V. finalized the manuscript. All the authors read and approved the final manuscript. F.X.C.V., S.W.I.S., and J.Y. acquired funding. The authors declare no conflict of interest.

## 6 Acknowledgments

The authors acknowledge the support of the Government of Canada’s New Frontiers in Research Fund (NFRF) (NFRFE-2021-00913), the Postdoctoral Fellowship Program of CPSF under Grant GZC20233322, and the Postdoctoral Talent Special Program. J.Y. was supported by the NFRF, the CPSF, and the Postdoctoral Talent Special Program. J.C. was the recipient of the Macau Polytechnic University graduate scholarship. The funders had no role in study design, data collection, and interpretation, or the decision to submit the work for publication.

## Data Availability

All peptide data used in this study comes from DBAASP [23] (https://dbaasp.org/home). The data and data preparation scripts for reproducing the experiments can be downloaded from https://github.com/jieluyan/Diffusion-ACP. The final prediction models trained in this study are also available in the previous link. The web server to directly access this prediction method is at https://app.cbbio.online/ACPep/home.

